# Comparing EEG/MEG neuroimaging methods based on localization error, false positive activity, and false positive connectivity

**DOI:** 10.1101/269753

**Authors:** Roberto D. Pascual-Marqui, Pascal Faber, Toshihiko Kinoshita, Kieko Kochi, Patricia Milz, Keiichiro Nishida, Masafumi Yoshimura

**Author notes:** Corresponding author: RD Pascual-Marqui; www.uzh.ch/keyinst/loreta.htm, scholar.google.com/citations?user=pascualmarqui.

## Abstract

EEG/MEG neuroimaging consists of estimating the cortical distribution of time varying signals of electric neuronal activity, for the study of functional localization and connectivity. Currently, many different imaging methods are being used, with very different capabilities of correct localization of activity and of correct localization of connectivity. The aim here is to provide a guideline for choosing the best (i.e. least bad) imaging method. This first study is limited to the comparison of the following methods for EEG signals: sLORETA and eLORETA (standardized and exact low resolution electromagnetic tomography), MNE (minimum norm estimate), dSPM (dynamic statistical parametric mapping), and LCMVBs (linearly constrained minimum variance beamformers). These methods are linear, except for the LCMVBs that make use of the quadratic EEG covariances. To achieve a fair comparison, it is assumed here that the generators are independent and widely distributed (i.e. not few in number), giving a well-defined theoretical population EEG covariance matrix for use with the LCMVBs. Measures of localization error, false positive activity, and false positive connectivity are defined and computed under ideal no-noise conditions. It is empirically shown with extensive simulations that: (1) MNE, dSPM, and all LCMVBs are in general incapable of correct localization, while sLORETA and eLORETA have exact (zero-error) localization; (2) the brain volume with false positive activity is significantly larger for MN, dSPM, and all LCMVBs, as compared to sLORETA and eLORETA; and (3) the number of false positive connections is significantly larger for MN, dSPM, all LCMVBs, and sLORETA, as compared to eLORETA. Non-vague and fully detailed equations are given. PASCAL program codes and data files are available. It is noted that the results reported here do not apply to the LCMVBs based on EEG covariance matrices generated from extremely few generators, such as only one or two independent point sources.

## 2. Introduction

This paper deals with the EEG neuroimaging problem: given non-invasive measurements of scalp electric potential differences, find the generators, in the form of time varying cortical distribution of electric neuronal activity. Focused reviews can be found in (Valdes-Sosa et al 2009; Pascual-Marqui 2009; Pascual-Marqui et al 2009).

There are available many different solutions, and the aim here is to provide a guideline for choosing the best (i.e. least bad) imaging method.

This first study is limited to the comparison of the following methods for EEG signals: 
- sLORETA and eLORETA (standardized and exact low resolution electromagnetic tomography) (Pascual-Marqui 2002; Pascual-Marqui 2007; Pascual-Marqui et al 2011).
- MNE (minimum norm estimate) (Hamalainen and Ilmoniemi 1994),
- dSPM (dynamic statistical parametric mapping) (Dale et al 2000),
- LCMVBs (linearly constrained minimum variance beamformers), consisting of three main variants, denoted as unit gain (UG), unit array gain (UAG), and unit noise gain (UNG); see e.g. (Van Veen et al 1997; Sekihara and Nagarajan 2008).

All these methods are linear, except for the LCMVBs, which make use of the EEG covariance, thus depending quadratically on the measurements. To achieve a fair comparison, it is assumed here that the generators are independent and distributed (i.e. not few in number), giving a well-defined theoretical population covariance matrix for the measurements to be used in the LCMVBs.

Measures of localization error, false positive activity, and false positive connectivity are defined and computed under ideal no-noise conditions.

Non-vague and fully detailed equations are given. PASCAL program codes and data files are available, allowing the interested researcher to check, test, validate, and replicate all the comparisons.

## 3. The forward EEG equation for the current density vector field

The general theory and formulation of the EEG/MEG forward problem can be found in e.g. (Sarvas 1987).

The discrete forward EEG equation, for the average reference, can be expressed as: 
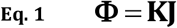
where 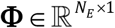 is the average reference scalp electric potential at *N*_*E*_ electrodes, 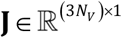 is the discrete current density vector field at *N*_*V*_ voxels in cortical grey matter, and 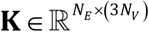 is the average reference lead field. The justification for using the average reference is explained in detail in e.g. (Pascual-Marqui et al 2011).

Note that at the i-th voxel, the current density vector is denoted as: 
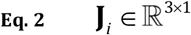
with: 
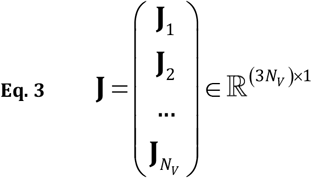

Note that at the i-th voxel, the lead field is denoted as: 
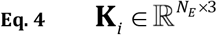
with: 
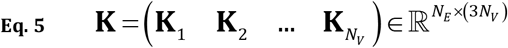

Note that average reference implies that: 
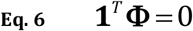
and: 
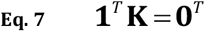
where 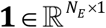 is a vector of ones, 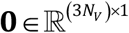 is a vector of zeros, and where the superscript “*T*” denotes vector-matrix transposed.

## 4. MNE (minimum norm estimate)

The minimum norm estimate (see e.g. Hamalainen and Ilmoniemi 1994) for the full current density distribution is: 
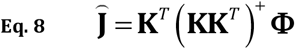
where the superscript “+” denotes the Moore-Penrose pseudoinverse. From here, the minimum norm inverse solution at the i-th voxel is given as: 
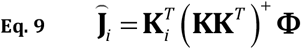

## 5. sLORETA (standardized low resolution electromagnetic tomography)

It will be assumed that the current densities are all independent and with equal variance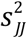 i.e.: 
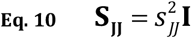
where 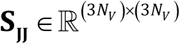 is the full current density covariance matrix, and **I** is the identity matrix.

From the forward equation Eq. 1, it follows that the EEG covariance is: 
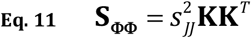

From the MNE (Eq. 8), it follows that its covariance is: 
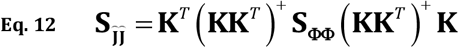
and plugging Eq. 11 into Eq. 12 gives: 
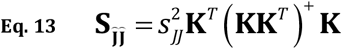

Thus, from Eq. 13, the variance 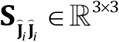 of the MNE at the i-th voxel is: 
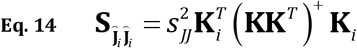

sLORETA at the i-th voxel is defined as the standardized MNE: 
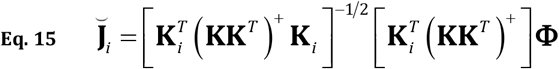

where the superscript “−1/2” denotes the symmetric square root inverse matrix, and where the theoretical variance at each voxel is set to one, i.e.: 
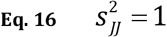

## 6. eLORETA (exact low resolution electromagnetic tomography)

eLORETA is a particular form of weighted minimum norm solution that, unlike all other weighted tomographies, uniquely achieves exact (zero-error) localization (Pascual-Marqui 2007; Pascual-Marqui et al 2011).

The full current density is: 
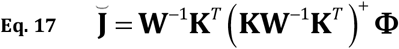
where the weight matrix 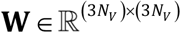 has all elements equal to zero, except for the diagonal subblocks denoted as **w**_*i*_ ∈ ℝ^3×3^ for each voxel *i* = 1…*N*_*V*_.

eLORETA at the i-th voxel is: 
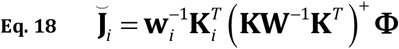

The eLORETA weights satisfy the following system of equations: 
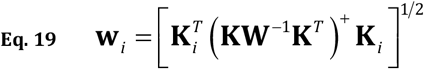
where the superscript “1/2” denotes the symmetric square root matrix.

## 7. dSPM (dynamic statistical parametric mapping)

The dSPM estimate at the i-th voxel is: 
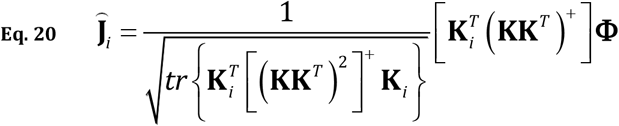
where the operator “*tr*” returns the trace of a matrix. Its detailed definition and derivation can be found in Dale et al (2000), and in Sekihara and Nagarajan (2008, see Equation 3.24 therein).

## 8. LCMVBs (linearly constrained minimum variance beamformers)

There are three popular beamformers, depending on the constraint type. For details, see e.g. (Van Veen et al 1997; Sekihara and Nagarajan 2008).

Note that in all cases below it will be assumed that that EEG covariance matrix is not estimated. In its place, a population EEG covariance will be used, under the assumption that the current densities are all independent and with equal variances, as used above for sLORETA, expressed in Eq. 10 and Eq. 11.

### UG-LCMVB (unit gain linearly constrained minimum variance beamformer)

The UG-LCMVB at the i-th voxel is: 
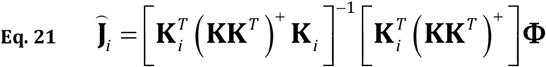

Its detailed definition and derivation can be found in (Van Veen et al 1997).

### UAG-LCMVB (unit array gain linearly constrained minimum variance beamformer)

The UAG-LCMVB at the i-th voxel is: 
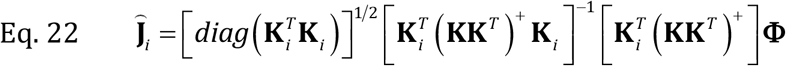
where the *diag*(•) operator sets all off-diagonal elements of a matrix to zero. The detailed definition and derivation of this particular version can be found in e.g. (Sekihara and Nagarajan 2008, see Equation 4.67 therein).

### UNG-LCMVB (unit noise gain linearly constrained minimum variance beamformer)

The UNG-LCMVB at the i-th voxel is: 
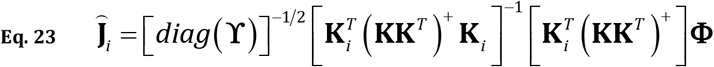

with: 
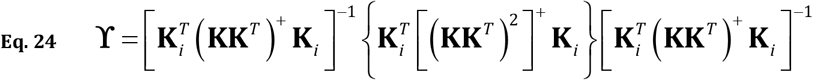

Its detailed definition and derivation can be found in e.g. (Sekihara and Nagarajan 2008, see Equation 4.85 therein).

## 9. Definition for “localization error”

Informally, the localization error is defined as the distance between the location of an actual localized source, and the location of the maximum of the estimated squared amplitude current density. The formal definition follows.

Let the actual point source be located at the *τ*-th voxel, and let ∣**Ĵ**_*i*_(τ)∣^2^ denote the estimated activity (i.e. current density squared amplitude) at the i-th voxel, due to the actual source at the τ-th voxel.

Let κ(τ) denote the voxel index at which the squared amplitude of the estimated current density attains its maximum value, i.e.: 
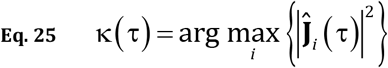

Note that κ(τ) will have an integer value in the range 1…*N*_*V*_.

Then the localization error in units of distance is defined as: 
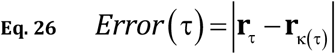
where in general **r**_*j*_ ∈ ℝ^3×1^ denotes the position vector of the j-th voxel.

## 10. Definition for “false positive activity”

Informally, false positive activity consists of the set of voxels with estimated activity higher than a certain fraction of the estimated activity at the location of the actual point source. The formal definition follows.

As before, let the actual point source be located at the τ-th voxel, and let ∣**Ĵ**_*i*_(τ)∣^2^ denote the estimated activity at the i-th voxel, due to the actual source at the τ-th voxel.

Given an actual generator at the τ-th voxel, the number of voxels with false positive activity is defined as: 
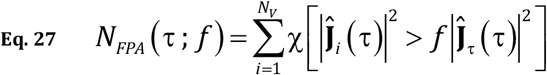
for a given fraction “*f*”, with 0 < *f* ≤ 1, and where “χ” is the indicator function, defined as: 
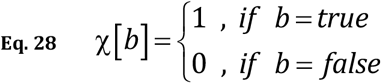

Finally, false positive activity is defined as the percent relative to the total number of voxels: 
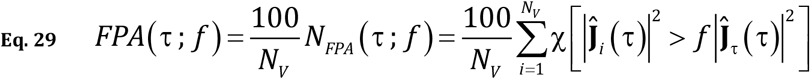

Note that false positive activity, as defined here, is also a measure of the spatial dispersion of the imaging method: a high value for the false positive activity indicates high spatial spreading of the imaging method.

Indeed, when for instance *f* = 0.5, the measure of false positive activity in Eq. 29 can be described as the “false active volume at half true source activity”.

## 11. Definition for “false positive connectivity”

In brain functional connectivity analysis, “connectivity” is commonly quantified by the simple “correlation” coefficient between a pair of signals measured at two cortical locations (see e.g. Worsley et al 2005).

Consider that ideal case where all cortical generators are truly independent. Then the estimated squared correlation coefficient between a pair of signals obtained with an EEG neuroimaging method should be small, below a certain threshold. A high value for the estimated squared correlation coefficient corresponds to a false positive connection. The formal definition for “false positive connectivity” follows.

It will be assumed that the current densities are all independent and with equal variances, as used above for sLORETA, and as expressed in Eq. 10 and Eq. 11.

Let the τ-th voxel be defined as the target voxel, and let *r̂*^2^(τ,*i*) denote the estimated squared correlation coefficient (based on a specific neuroimaging method) between the target voxel and all others, for *i* = 1…*N*_*v*_ and *i* ≠ τ. Note that *r̂*^2^(τ,*i*) satisfies: 0≤*r̂*^2^(τ,*i*)≤1.

Given a threshold 0 < ρ < 1, a connection between the τ-th target voxel and the i-th voxel is defined as a “false positive connection” if: 
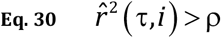

For the target voxel “τ”, and for the threshold value ρ, the number of false positive connections is: 
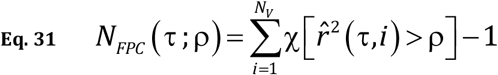
where “χ” is the indicator function defined in Eq. 28.

Finally, false positive connectivity is defined as the percent relative to the total number of connections with the τ-th voxel: 
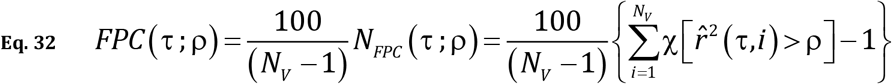

It now remains to define the estimator for the squared correlation between voxels, under the assumption of true total independence.

Note that all the neuroimaging methods studied here (Eq. 9, Eq. 15, Eq. 18, Eq. 20, Eq. 21, Eq. 22, Eq. 23) can be expressed as: 
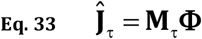
where **Ĵ**_τ_ denotes the estimated current density at the τ-th voxel, and where 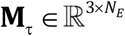 is the linear inverse operator specific to each neuroimaging method.

Under the assumption that the current densities are all independent and with equal variances, as used above for sLORETA, expressed in Eq. 10 and Eq. 11, the matrix of correlation coefficients **R̂**_*τi*_ ∈ ℝ^3×3^ between the τ-th and i-th voxels is: 
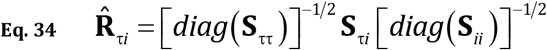
with covariance matrices: 
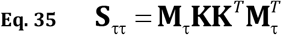

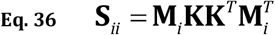

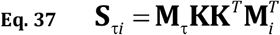

From the nine correlation coefficients in **R̂**_*τi*_ ∈ ℝ^3×3^ (Eq. 34), compute the corresponding nine squared correlations, and define *r̂*^2^(τ, *i*) (used in Eq. 30, Eq. 31, Eq. 32) as the maximum of the nine values.

## 12. The toy data: head model, voxels, electrodes

A unit radius three-shell spherical head model was used, as described in (Ary et al 1981). The number of voxels was *N*_*V*_=818, corresponding to a regular 3D grid at 0.133 units resolution, with maximum radius=0.8, and with vertical coordinate values *Z* ≥−0.4. The number of scalp electrodes was *N*_*E*_ = 148, uniformly distributed over the outer sphere surface for
*Z* ≥−0.5. The lead field 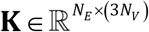 was computed for this simple head model using the approximations by (Ary et al 1981).

## 13. Results

### Localization error

For each imaging method, the localization errors in Eq. 26 were computed, for all possible 2454 actual generators, corresponding to τ = 1…818 voxels, and three dipole moments. The localization errors as a function of the actual source radius are shown in Figure 1a.

**Figure 1a:**
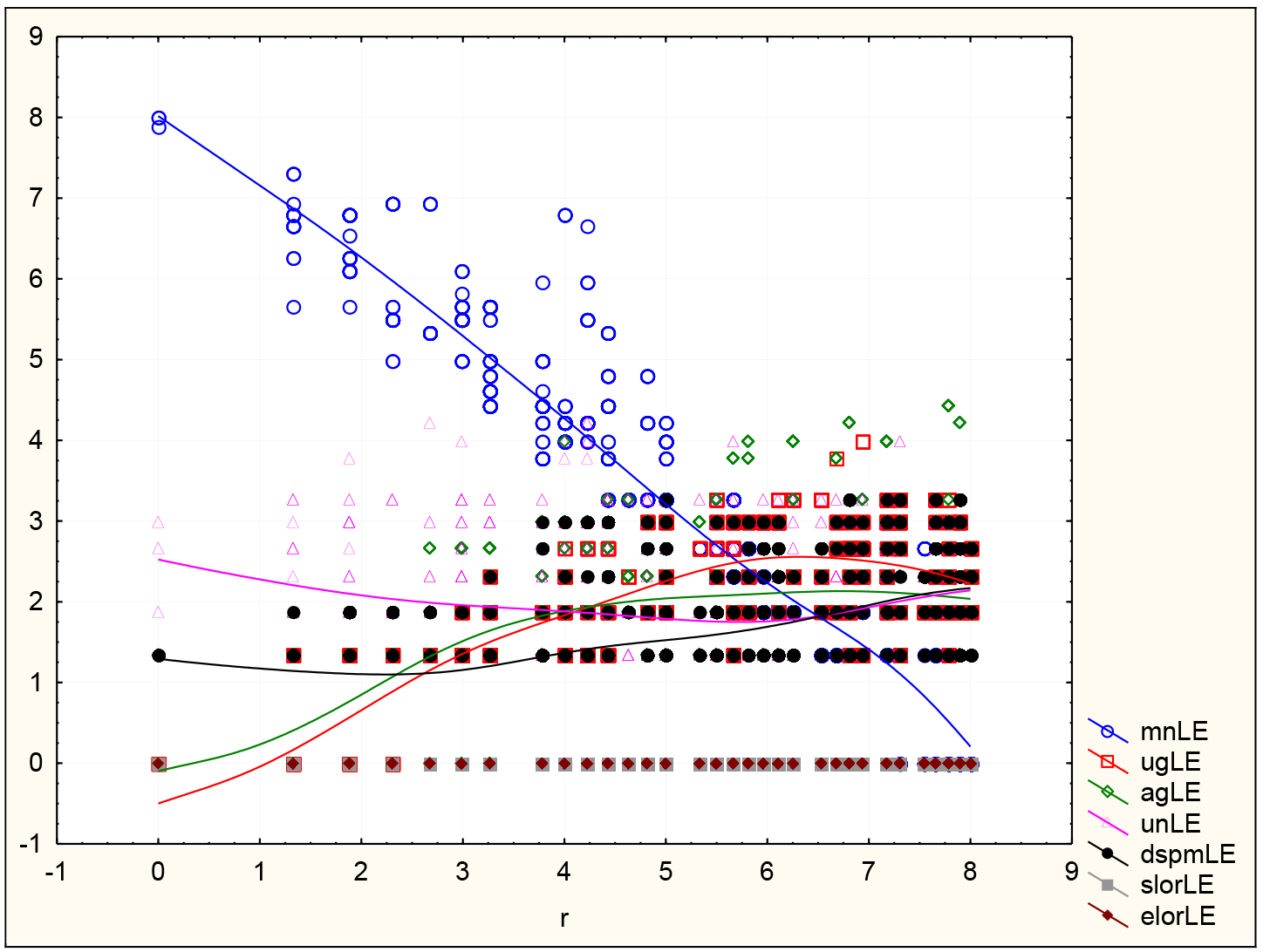
For each of seven neuroimaging methods, the localization errors in Eq. 26 were computed, for all possible 2454 actual generators, corresponding to τ = 1…818 voxels, and three dipole moments. Vertical axis: localization error in mm. Horizontal axis: radius of the actual source in cm, assuming a 10 cm radius head. mn: minimum norm; ug: unit gain beamformer; ag: array gain beamformer; un: unit noise gain beamformer; dspm: dynamic SPM; slor: sLORETA; elor: eLORETA.

Figure 1b displays a box-and-whiskers plot for the mean localization error, its standard error, and 1.96 times its standard error.

**Figure 1b:**
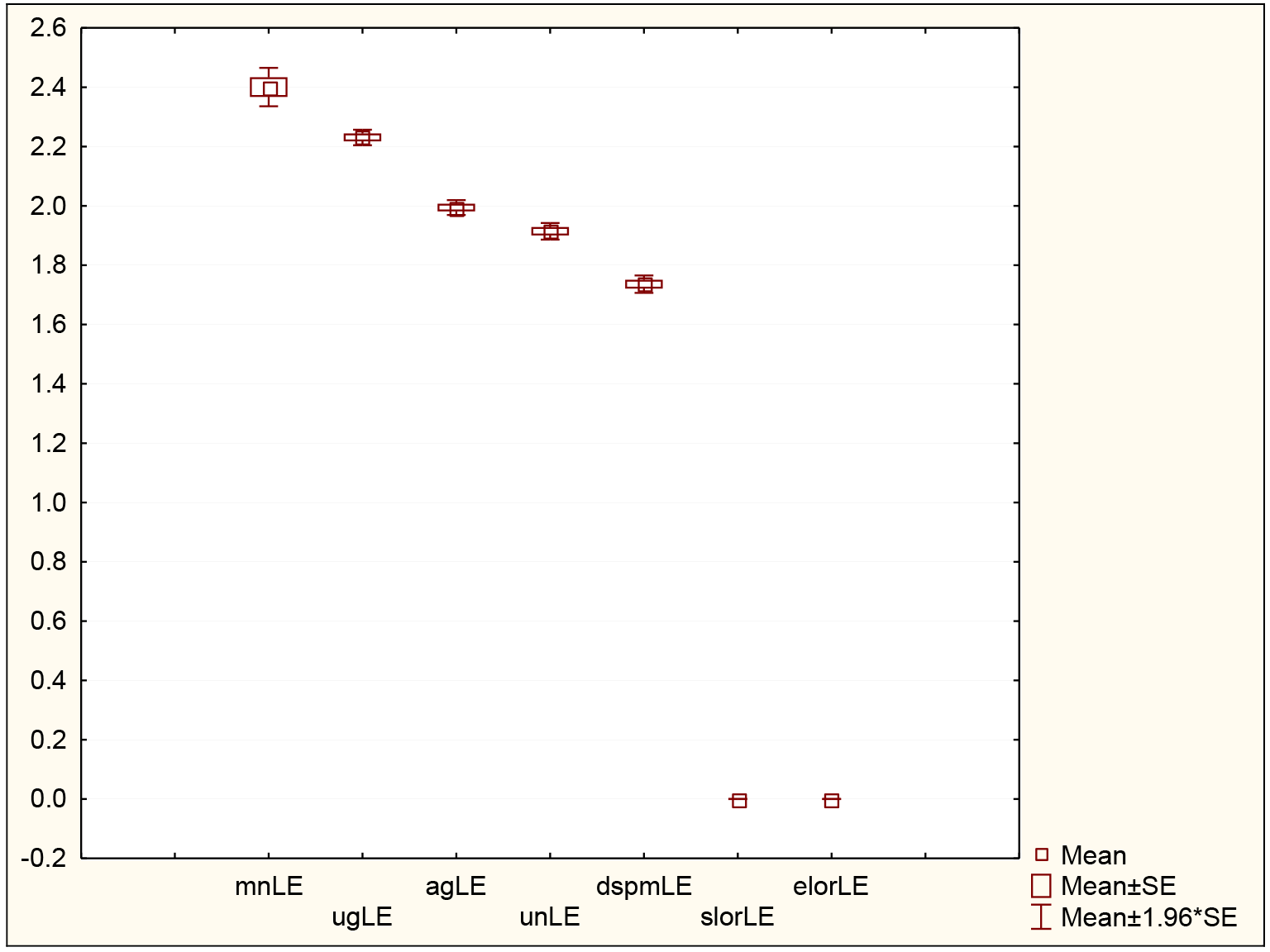
For each of seven neuroimaging methods, a box-and-whiskers plot for the mean localization error, its standard error, and 1.96 times its standard error. These statistics are computed for all possible 2454 actual generators, corresponding to τ = 1…818 voxels, and three dipole moments. Vertical axis: localization error in mm. mn: minimum norm; ug: unit gain beamformer; ag: array gain beamformer; un: unit noise gain beamformer; dspm: dynamic SPM; slor: sLORETA; elor: eLORETA.

Table 1 shows the t-tests comparing all pairs of neuroimaging methods based on localization error.

**Table 1:** Comparison of imaging methods based on localization error

|  | mnLE | ugLE | agLE | unLE | dspmLE | slorLE | elorLE |
| --- | --- | --- | --- | --- | --- | --- | --- |
| mnLE | 0.0 | 4.1 | 10.5 | 13.5 | 16.4 | 72.7 | 72.7 |
| ugLE | -4.1 | 0.0 | 18.7 | 17.0 | 30.8 | 167.1 | 167.1 |
| agLE | -10.5 | -18.7 | 0.0 | 4.2 | 14.5 | 155.4 | 155.4 |
| unLE | -13.5 | -17.0 | -4.2 | 0.0 | 10.6 | 134.2 | 134.2 |
| dspmLE | -16.4 | -30.8 | -14.5 | -10.6 | 0.0 | 115.9 | 115.9 |
| slorLE | -72.7 | -167.1 | -155.4 | -134.2 | -115.9 | 0.0 | 0.0 |
| elorLE | -72.7 | -167.1 | -155.4 | -134.2 | -115.9 | 0.0 | 0.0 |

T-tests (row variable minus column variable) comparing all pairs of neuroimaging methods based on localization error. Red and green highlight indicate significant t-values at $p < 10^{-6}$ . For instance, the last two rows corresponding to sLORETA and eLORETA with extreme negative t-values demonstrate that all other imaging methods have significantly larger localization errors. In fact, sLORETA and eLORETA have exactly zero error localization. mn: minimum norm; ug: unit gain beamformer; ag: array gain beamformer; un: unit noise gain beamformer; dspm: dynamic SPM; slor: sLORETA; elor: eLORETA.

### False positive activity

For each imaging method, false positive activity in Eq. 29 was computed, for all possible 2454 actual generators, corresponding to τ = 1…818 voxels, and three dipole moments. The fraction value (*f*=0.5) of the actual generator amplitude was used. Figure 2a and Figure 2b display false positive activity as a function of the actual source radius.

**Figure 2a:**
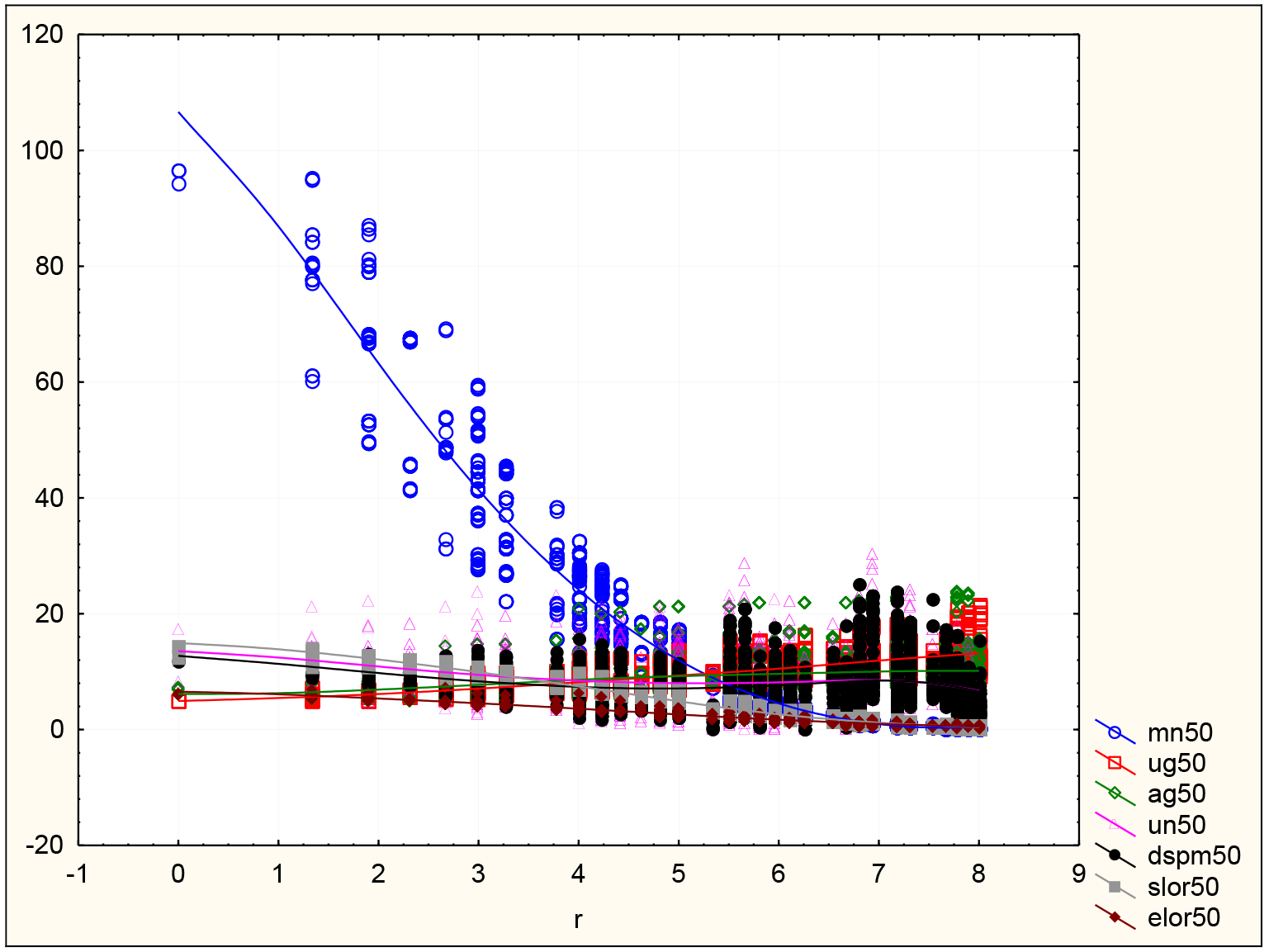
For each of seven neuroimaging methods, the false positive activity in Eq. 29 was computed, for all possible 2454 actual generators, corresponding to τ =1…818 voxels, and three dipole moments. Vertical axis: false positive activity, at the fraction (*f*=0.5) of the actual generator amplitude, as percent of the total number of actual sources. Horizontal axis: radius of the actual source in cm, assuming a 10 cm radius head. mn: minimum norm; ug: unit gain beamformer; ag: array gain beamformer; un: unit noise gain beamformer; dspm: dynamic SPM; slor: sLORETA; elor: eLORETA.

In Figure 2a, the MNE shows extremely high false positive activation for actual deep sources. The detailed behavior of all other imaging methods cannot be appreciated. Figure 2b excludes the MNE.

**Figure 2b:**
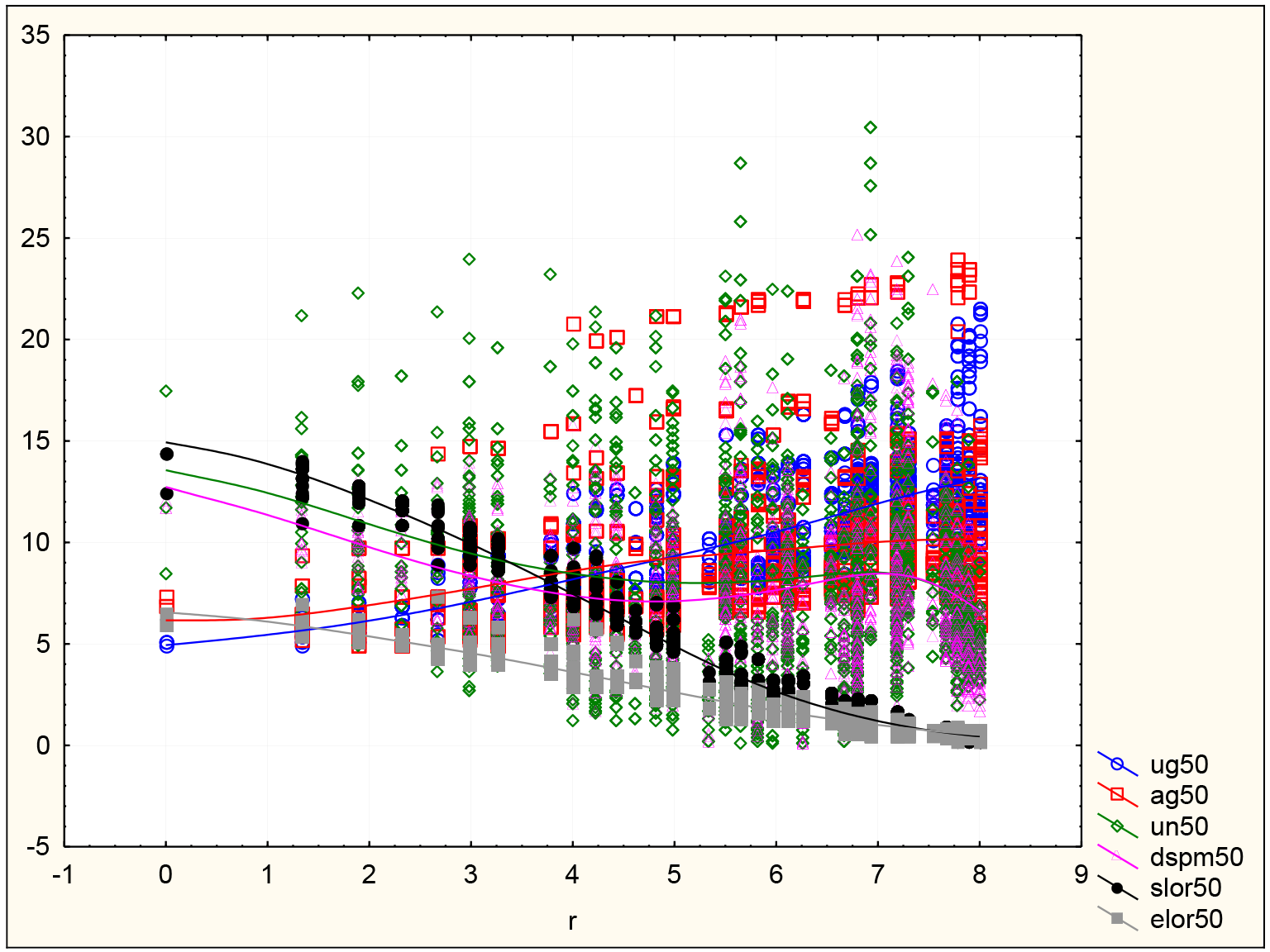
For each of six neuroimaging methods, the false positive activity in Eq. 29 was computed,
for all possible 2454 actual generators, corresponding to τ =1…818 voxels, and three dipole moments. Vertical axis: false positive activity, at the fraction (*f*=0.5) of the actual generator amplitude, as percent of the total number of actual sources. Horizontal axis: radius of the actual source in cm, assuming a 10 cm radius head. ug: unit gain beamformer; ag: array gain beamformer; un: unit noise gain beamformer; dspm: dynamic SPM; slor: sLORETA; elor: eLORETA.

Figure 2c displays a box-and-whiskers plot for the mean false positive activity at the fraction (*f*=0.5), its standard error, and 1.96 times its standard error.

**Figure 2c:**
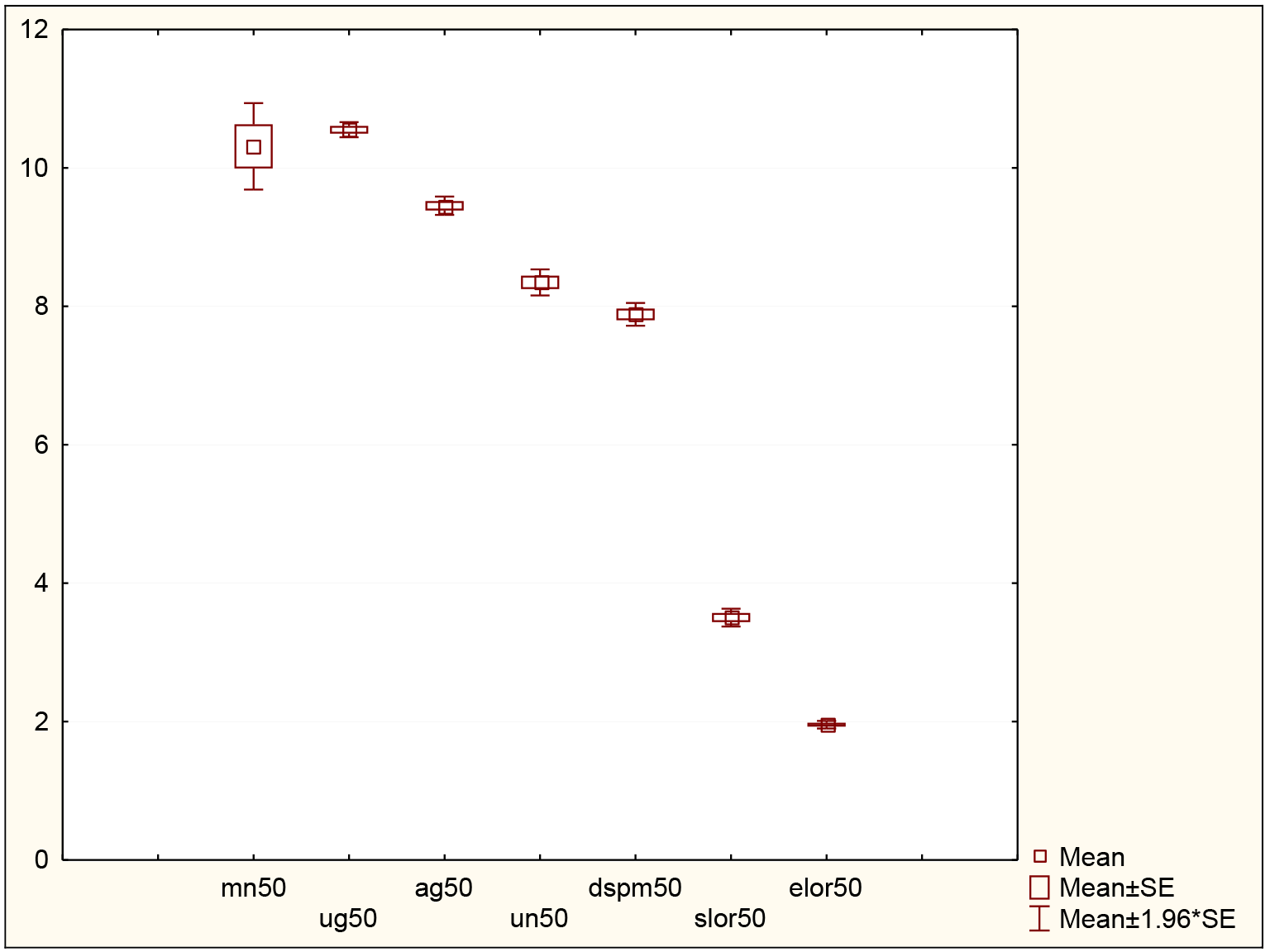
For each of seven neuroimaging methods, a box-and-whiskers plot for the mean false positive activity at the fraction (*f*=0.5), its standard error, and 1.96 times its standard error. These statistics are computed for all possible 2454 actual generators, corresponding to τ =1…818 voxels, and three dipole moments. Vertical axis: false positive activity, at the fraction (*f*=0.5) of the actual generator amplitude, as percent of the total number of actual sources. mn: minimum norm; ug: unit gain beamformer; ag: array gain beamformer; un: unit noise gain beamformer; dspm: dynamic SPM; slor: sLORETA; elor: eLORETA.

Table 2 shows the t-tests comparing all pairs of neuroimaging methods based on false positive activity.

**Table 2:** Comparison of imaging methods based on false positive activity

|  | mn50 | ug50 | ag50 | un50 | dspm50 | slor50 | elor50 |
| --- | --- | --- | --- | --- | --- | --- | --- |
| mn50 | 0.0 | -0.7 | 2.5 | 6.1 | 7.5 | 26.2 | 28.3 |
| ug50 | 0.7 | 0.0 | 25.0 | 21.2 | 28.8 | 64.2 | 113.4 |
| ag50 | -2.5 | -25.0 | 0.0 | 10.6 | 16.2 | 58.6 | 99.0 |
| un50 | -6.1 | -21.2 | -10.6 | 0.0 | 5.5 | 43.7 | 65.0 |
| dspm50 | -7.5 | -28.8 | -16.2 | -5.5 | 0.0 | 41.0 | 66.0 |
| slor50 | -26.2 | -64.2 | -58.6 | -43.7 | -41.0 | 0.0 | 39.8 |
| elor50 | -28.3 | -113.4 | -99.0 | -65.0 | -66.0 | -39.8 | 0.0 |

T-tests (row variable minus column variable) comparing all pairs of neuroimaging methods based on false positive activity. Red and green highlight indicate significant t-values at $p < 10^{-6}$ . For instance, the last two rows corresponding to sLORETA and eLORETA with extreme negative t-values demonstrate that all other imaging methods have significantly larger rates of false positive activity, with eLORETA better than sLORETA. mn: minimum norm; ug: unit gain beamformer; ag: array gain beamformer; un: unit noise gain beamformer; dspm: dynamic SPM; slor: sLORETA; elor: eLORETA.

### False positive connectivity

For each imaging method, false positive connectivity in Eq. 32, for each target voxel “τ” (for ((τ =1…818), was computed. A relatively high threshold value of (ρ=0.5) was used for declaring a connection as false. Figure 3a shows false positive connectivity as a function of the target voxel radius.

**Figure 3a:**
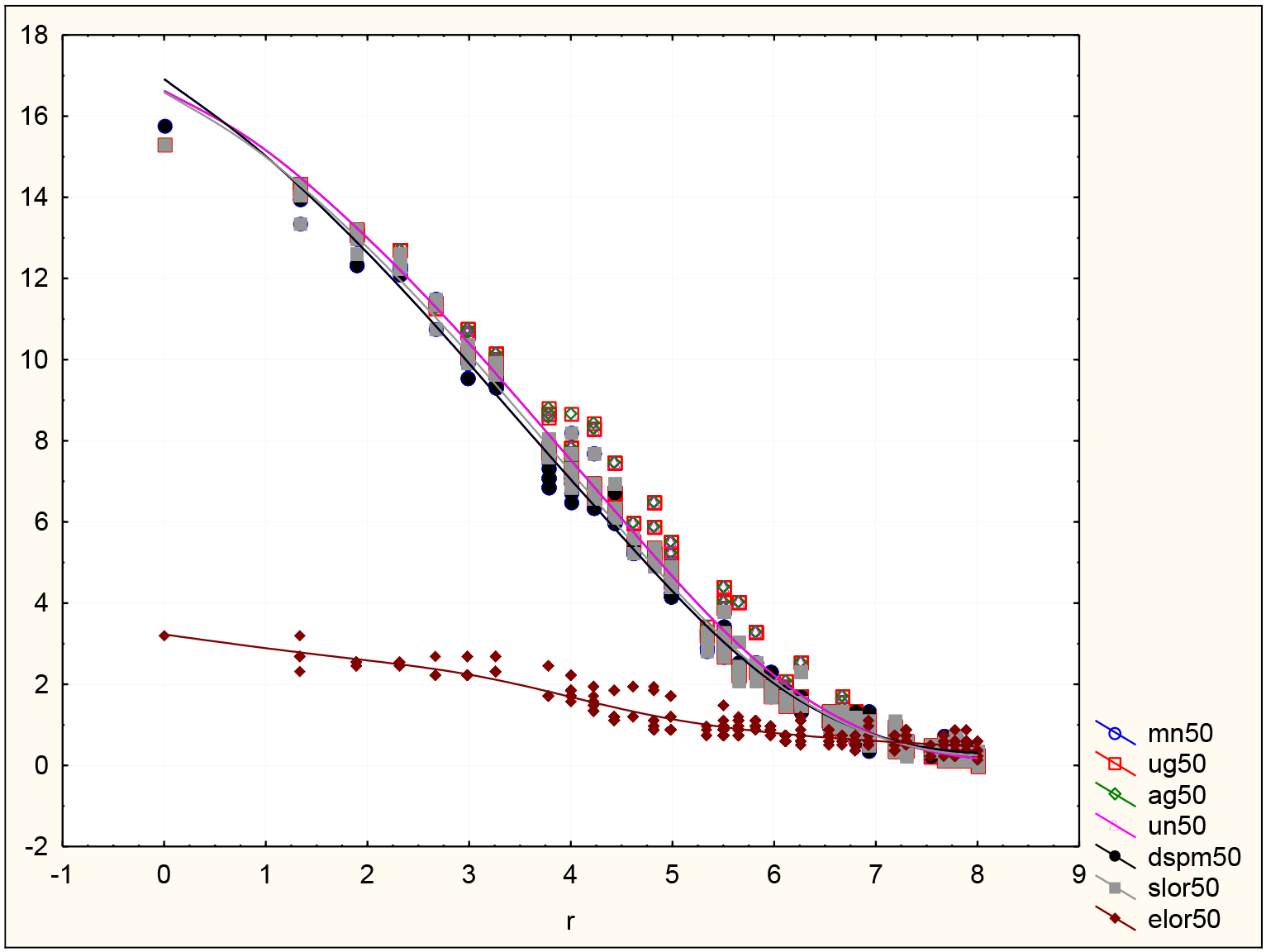
For each of seven neuroimaging methods, the false positive connectivity in Eq. 32 was computed at each target voxel (818 voxels). Vertical axis: false positive connectivity at threshold (ρ=0.5), as percent of the total number of connections with the target voxel. Horizontal axis: radius of the actual source in cm, assuming a 10 cm radius head. mn: minimum norm; ug: unit gain beamformer; ag: array gain beamformer; un: unit noise gain beamformer; dspm: dynamic SPM; slor: sLORETA; elor: eLORETA.

Figure 3b displays a box-and-whiskers plot for the mean false positive connectivity at threshold (ρ=0.5), its standard error, and 1.96 times its standard error.

**Figure 3b:**
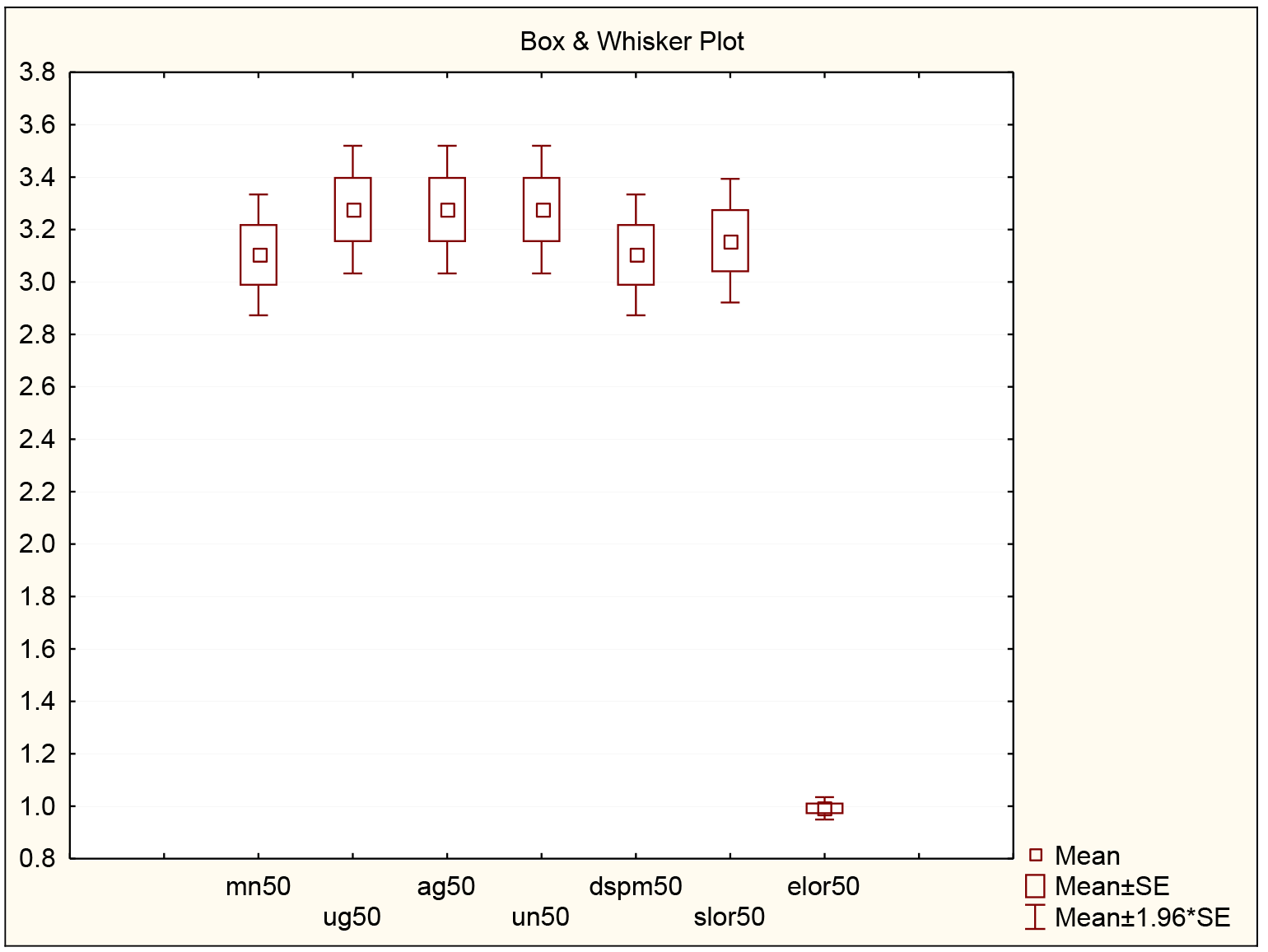
For each of seven neuroimaging methods, a box-and-whiskers plot for the mean false positive connectivity at threshold (ρ=0.5), its standard error, and 1.96 times its standard error. These statistics are computed for all possible 818 target voxels. Vertical axis: false positive connectivity at threshold (ρ=0.5), as percent of the total number of connections with the target voxel. mn: minimum norm; ug: unit gain beamformer; ag: array gain beamformer; un: unit noise gain beamformer; dspm: dynamic SPM; slor: sLORETA; elor: eLORETA.

Table 3 shows the t-tests comparing all pairs of neuroimaging methods based on false positive connectivity.

**Table 3:** Comparison of imaging methods based on false positive connectivity

|  | mn50 | ug50 | ag50 | un50 | dspm50 | slor50 | elor50 |
| --- | --- | --- | --- | --- | --- | --- | --- |
| mn50 | 0.0 | -13.4 | -13.4 | -13.4 | 0.0 | -7.4 | 21.6 |
| ug50 | 13.4 | 0.0 | 0.0 | 0.0 | 13.4 | 14.4 | 22.0 |
| ag50 | 13.4 | 0.0 | 0.0 | 0.0 | 13.4 | 14.4 | 22.0 |
| un50 | 13.4 | 0.0 | 0.0 | 0.0 | 13.4 | 14.4 | 22.0 |
| dspm50 | 0.0 | -13.4 | -13.4 | -13.4 | 0.0 | -7.4 | 21.6 |
| slor50 | 7.4 | -14.4 | -14.4 | -14.4 | 7.4 | 0.0 | 21.6 |
| elor50 | -21.6 | -22.0 | -22.0 | -22.0 | -21.6 | -21.6 | 0.0 |

T-tests (row variable minus column variable) comparing all pairs of neuroimaging methods based on false positive connectivity. Red and green highlight indicate significant t-values at $p<10^{-6}$ . For instance, the last row corresponding to eLORETA with extreme negative t-values demonstrates that all other imaging methods have significantly larger rates of false positive connections. mn: minimum norm; ug: unit gain beamformer; ag: array gain beamformer; un: unit noise gain beamformer; dspm: dynamic SPM; slor: sLORETA; elor: eLORETA.

## 14. Discussion

### Localization error

From a strict point of view, localization error is unacceptable in functional imaging, where one of the main goals is to correctly localize brain function. In other words: if a method with localization error is selected for neuroimaging studies, then it is almost certain that reported locations of brain function are incorrect.

Under the ideal testing conditions presented here, only sLORETA and eLORETA are capable of exact localization, with all other methods suffering from incapability of correct localization.

From best to worst, the ranking of neuroimaging methods based on localization error is: 
1. eLORETA, sLORETA
2. dSPM, all LCMVBs
3. MNE

### False positive activity

By far, the MNE method has extremely high values of false positive activity, especially when the actual generator is deep. The reason for this is because the MNE has maximum activity at the outermost voxels, even when the actual generator is deep. False positive activity, as defined here, is dependent on the estimated activity value at the actual generator location, and not at the very incorrect location of the MNE maximum activity.

Another interpretation follows: Note that false positive activity as defined here is very closely related to spatial spreading, or blurring. And note that spatial spreading is defined by computing the “spread”, not centered at the maximum location (which can be completely wrong), but centered at the actual generator location. This explains why the MNE has such high “spatial spreading”.

Under the ideal testing conditions presented here, eLORETA has an average false positive activity rate (at half maximum activity) of only 2%, with sLORETA at 3.5%, and all other methods ranging from 8% to 10.5%.

From best to worst, the ranking of neuroimaging methods based on localization error is: 
1. eLORETA
2. sLORETA
3. dSPM, all LCMVBs
4. MNE

### False positive connectivity

It is a well-known fact that all linear inverse solutions have low resolution, as can be confirmed by the results of “false positive activity” previously discussed. This means that in the best of cases, the neuroimaging-based estimated signal at any cortical location corresponds to an instantaneous linear mixture of all true cortical signals. Therefore, truly independent cortical generators will appear to be correlated based on estimated signals from neuroimaging methods. These are “false positive connections”.

False positive connectivity, as defined here, provides a measure of the number of false connections that are due to the properties of the neuroimaging method.

Under the ideal testing conditions presented here, eLORETA is by far the best neuroimaging method with the lowest rate of false positive connectivity. At the relatively high threshold of ((ρ=0.5), eLORETA has an average of 1% of false positive connections, while all other methods have an average of 3.2% of false positive connections.

From best to worst, the ranking of neuroimaging methods based on false positive connectivity is: 
1. eLORETA
2. MNE, dSPM, sLORETA, all LCMVBs

It should be noted that there have been attempts in the literature aimed at “decorrelating” the estimated signals. One such recent attempt was developed by Colclough et al (2015). However, it was demonstrated in (Pascual-Marqui et al 2017) that the Colclough et al (2015) method actually produces false connectivity results. A new method, denoted “innovations orthogonalization” (Pascual-Marqui et al 2017), was proposed and validated, that properly decorrelates the estimated signals.

## 15. Conclusions

Of all tested methods, eLORETA excels in the three indices, having: 
1. Exact (zero-error) localization,
2. Minimum rate of false positive activity, which at the same time means it has the highest resolution (i.e. lowest spatial spreading), and,
3. Minimum rate of false positive connectivity.

The next best EEG neuroimaging method, based on these three indices, is sLORETA.

This is followed by the remaining methods (dSPM and all three LCMVBs), which perform poorly.

The worst method is the MNE.

Near future extended tests must (and will) include MEG, and noise, and connectivity measures of the Granger (causality) type, such as the isolated effective coherence “iCoh” (Pascual-Marqui et al 2014).

Meanwhile, it is worth emphasizing that the results presented here are in disagreement with those of Anzolin et al (2018), where they compare eLORETA with a LCMVB, and claim improved performance of the LCMVB.

Some other few recent papers have made some comparisons of beamformers and LORETA (Pascual-Marqui et al 1994) or sLORETA (Pascual-Marqui 2002), claiming superiority of the LCMVBs, see e.g. (Little et al 2018; Bonaiuto et al 2018). In those papers, it is not even clear which method was used (if LORETA of sLORETA). In any case, the results presented here challenge the superiority of the LCMVBs in the case of an EEG/MEG covariance matrix generated by distributed sources, and not only one or two independent sources.

